# Chimeric oncolytic adenovirus evades neutralizing antibodies from human patients and exhibits enhanced anti-glioma efficacy in immunized mice

**DOI:** 10.1101/2023.07.11.548552

**Authors:** Dong Ho Shin, Hong Jiang, Andrew Gillard, Debora Kim, Xuejun Fan, Sanjay Singh, Teresa T Nguyen, Sagar Sohoni, Andres Lopez-Rivas, Akhila Parthasarathy, Chibawanye I. Ene, Joy Gumin, Frederick Lang, Marta M Alonso, Candelaria Gomez-Manzano, Juan Fueyo

## Abstract

**BACKGROUND:** Oncolytic adenoviruses, such as Delta-24-RGD, show promise as a potential breakthrough in treating patients with high-grade gliomas. However, their effectiveness against gliomas can be hindered by the presence of neutralizing antibodies.

**METHODS:** Production of human neutralizing antibodies against adenoviruses was assessed in two cohorts of patients with malignant gliomas treated with Delta-24-RGD in a phase 1 clinical trial. Sera containing neutralizing antibodies were also obtained from mice immunized with intramuscular injections of wild-type Ad5. Chimeric adenovirus was constructed using molecular cloning, and its activity was assessed in vitro using quantitative PCR, western blot, and transmission electron microscopy. The therapeutic efficacy of the chimeric virus was tested in vivo using sera from patients previously treated with Delta-24-RGD and immunocompetent murine models of glioma.

**RESULTS:** Examination of sera from patients with malignant gliomas treated with Delta-24-RGD revealed that in the cohort treated with multiple injections of this oncolytic adenovirus, a higher percentage of patients developed neutralizing antibodies when compared to the patients treated with a single injection of Delta-24-RGD. Of note, long-term survival was only observed in patients who received a single injection. Delta-24-RGD-H43m, a chimeric oncolytic adenovirus engineered to overcome virus neutralization, demonstrated a potent anti-glioma effect both in vitro and in vivo. This chimeric virus showed resilience against anti-Ad5 neutralizing antibodies and conferred better therapeutic efficacy compared to Delta-24-RGD in mice with immunity against Ad5. Of further clinical relevance, Delta-24-RGD-H43m also evaded the inhibitory effects of sera from human patients treated with Delta-24-RGD.

**CONCLUSIONS:** The development of neutralizing antibodies due to multiple virus injections was associated with lower frequency of long-term survivors in a clinical trial. The new chimeric virus shows increased resilience to inactivation by the sera of human patients compared to the parental virus. These findings lay the foundation for a novel oncolytic virus treatment approach targeting a significant percentage of glioma patients with prior exposure to adenovirus.

## INTRODUCTION

We have developed a platform of oncolytic viruses (OV) based on the backbone of an Rb-targeted adenovirus termed Delta-24-RGD to treat brain tumors.^1,2^ While survival rates for many cancers have improved significantly over the past several years due to progress in early detection, the advent of precision medicine and advances in treatments, glioblastoma (GBM) continues to be an exception to this trend, with long-term survivorship remaining unacceptably low. Furthermore, immunotherapy with immune checkpoint inhibitors, which has been successful in other tumors, has not conferred a survival benefit in most patients with malignant gliomas, suggesting that these tumors are immunologically “cold” and, therefore, resistant to immune checkpoint antibodies.^3^ Clinical trials have indicated that oncolytic adenoviruses are a promising new category of anticancer biotherapeutic agents; in fact, the first oncolytic virus approved for standard clinical practice in 2006 was an adenovirus.^4^ A first-in-human phase I clinical trial of Delta-24-RGD was successfully completed in patients with recurrent GBMs (NCT00805376). In this trial, we observed durable (>3 years) responses in 20% of patients.^5^ In agreement with these data, similar results were confirmed in two additional clinical trials in patients with DIPG and recurrent GBM.^6,7^ Results from these trials suggest that Delta-24-RGD induces strong immune responses and durable tumor regression in a subset of patients, but what differentiates responders from non-responders remains unclear.

Interestingly, analyses of data from pediatric patients showed that adenovirus-specific neutralizing antibody (NAb) titers could be used to stratify patients by survival.^6^ Patients that developed higher-than-median NAb titers had a median survival of 12.5 months, whereas those with lower-than-median NAb titers had a median survival of 21.3 months. This discovery led us to examine the role of NAbs in determining the efficacy of OVs locally injected into brain tumors. Delta-24-RGD is built upon an adenovirus serotype 5 (Ad5) backbone. Ad5 is the most widely utilized adenoviral vector in cancer and gene therapies but is also among the most prevalent serotypes.^8^ These factors indicate that the development of NAbs can influence the outcomes of oncolytic virotherapies.

In this work, we showed that intratumoral virus injection leads to the development of NAbs in patients with malignant gliomas enrolled in a phase 1 clinical trial (NCT00805376),^5^ and we discovered the cohort with the highest percentage of patients developing NAbs received multiple OV injections and intriguingly exhibited poorer long-term survival benefits. Based on these data, we hypothesized that evasion of NAbs will improve the anti-glioma efficacy of virotherapy. To test this hypothesis, we generated a chimeric oncolytic adenovirus called Delta-24-RGD-H43m. We found that this chimeric OV evades anti-Ad5 NAbs presented in sera from mice and yields superior outcomes in orthotopic murine models of glioma. Of clinical interest, we also demonstrated that Delta-24-RGD-H43m evades NAbs from patients previously treated with Delta-24-RGD. These data provide the rationale for the development of clinical trials poised to examine the effect of NAb-resistant virotherapy in patients with gliomas and other solid tumors.

## RESULTS

### Intratumoral Delta-24-RGD injection leads to the development of NAbs in patients with malignant gliomas

To examine the generation of neutralizing antibodies (NAbs) after virotherapy, we analyzed sera from a completed phase 1 clinical trial (NCT00805376) involving 37 patients with recurrent malignant gliomas treated with Delta-24-RGD.^5^ In this trial, 37 patients were divided into two groups: twenty-five patients, in group A, received a single stereotactic injection of Delta-24-RGD into tumors and were followed for toxicity and clinical outcomes; twelve patients, in group B, received an intratumoral injection of Delta-24-RGD, followed two weeks after by en-bloc tumor resection and multiple injections of Delta-24-RGD in the surgical cavity.^5^ Sera were collected from these patients at various time points, including before the initial virus injection (baseline), 1 month and 4 months after virus injections (figure 1A). To assess the presence of NAbs, we infected A549 cells with Delta-24-RGD in the presence of sera from these patients at several concentrations and quantified cell viability 48 hours later as a surrogate of virus activity. When we calculated serum dilutions that inhibited more than 50% of virus activity compared to no serum addition, we observed that approximately 40% of the patients from both group A and group B developed NAbs one month after virus injection (figure 1B). Importantly, although the percentage of patients in group A with NAbs did not change at 4 months after treatment, 82% of the patients from group B developed NAbs (*p*=0.05, paired t-test; figure 1B, supplementary figure S1). This data immediately suggested that multiple virus injections received by patients enrolled in group B resulted in a higher percentage of patients developing NAbs. Interestingly, there were also differences in the overall survival of patients in groups A and B. Thus, no patients from group B survived more than 3 years, whereas 20% of the patients from group A, which has a significantly lower percentage of patients with NAbs, survived more than 3 years (figure 1C). This inverse relationship between NAb development and long-term survival led us to further examine the role of NAbs during glioma virotherapy.

**Figure 1.**
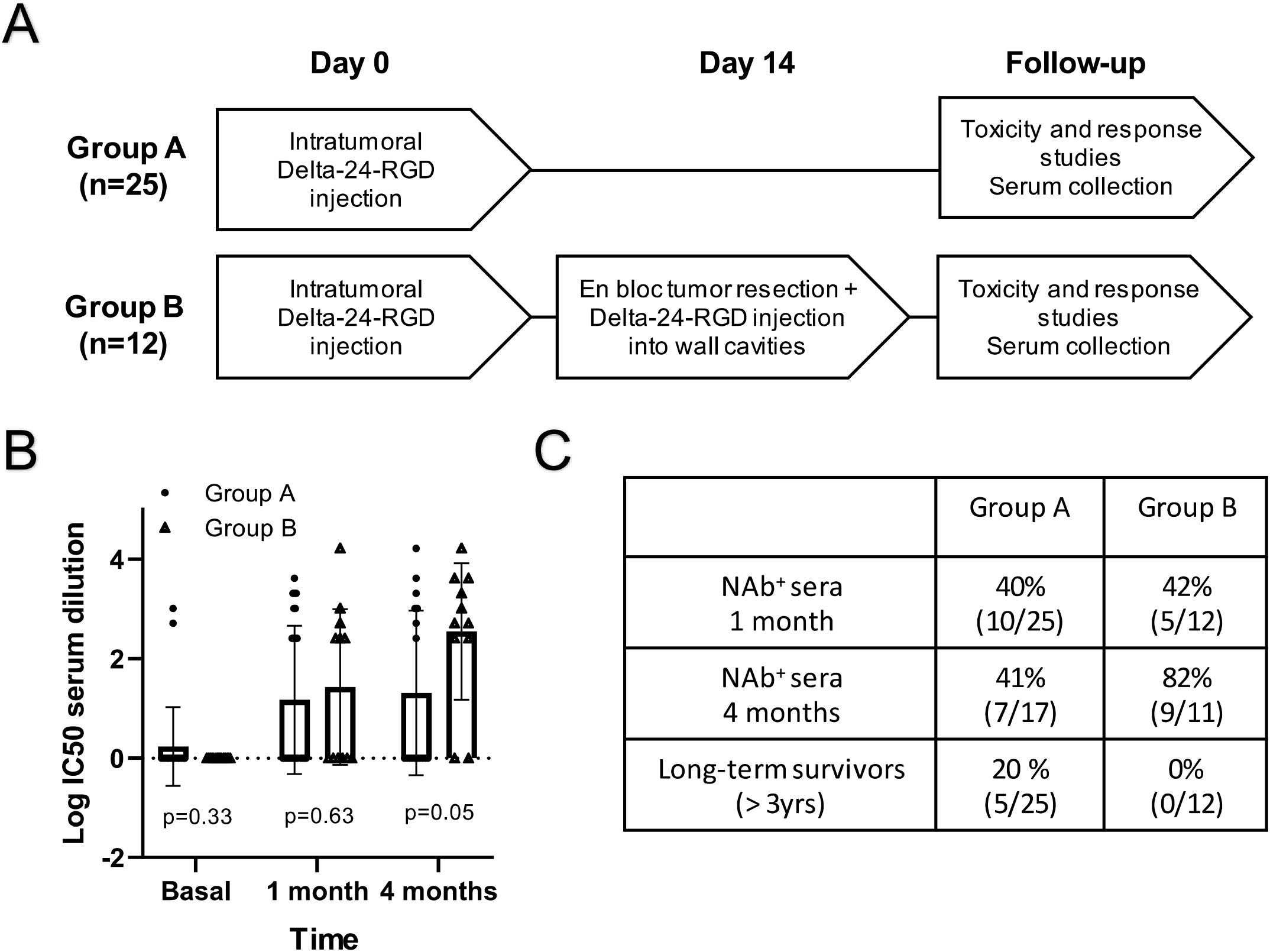
Patients with recurrent GBM developed NAbs after Delta-24-RGD treatment. (A) Patients with recurrent high-grade gliomas who were enrolled in a phase I trial (NCT00805376) received intratumoral injections of Delta-24-RGD (1 x 10 to 3 x 10 viral particles). In group A (n=25), the patients received a single intratumoral virus injection. In group B (n=12), the patients received an intratumoral virus injection, followed by *en bloc* tumor resection two weeks later and several doses of virus injection into the wall of the surgical cavities. Sera were collected at various time points. (B) A549 cells were infected with Delta-24-RGD in the presence of patient sera. Cell viability was measured 48 hours later to identify the serum dilutions that inhibit 50% of cell death. Each dot represents serum from each patient; the *p* value was calculated using a paired t-test. (C) Table showing percentage of NAb-positive sera from each group at 1 month and 4 months after virus injections, and percentage of long-term survivors.

### Virotherapy induces the influx of antibodies into brain tumors

To determine whether NAbs might be present in gliomas upon OV intratumoral injection, we assessed the presence of immunoglobulins in a murine orthotopic glioma model. We intratumorally injected Delta-24-RGD into GSC005 murine glioma^9^–bearing C57BL/6 mice previously immunized against Ad5. After performing transcardial perfusion with saline to clear blood from the brains, we assessed intratumoral viral replication and antibody presence by immunofluorescence staining of viral hexon and mouse immunoglobulin G (IgG), respectively. As expected, we detected hexon protein in the brains from virus-injected mice but not in those of mock-treated mice (*p*=0.007, unpaired t-test; figure 2A,B). Of interest, we detected approximately a 30% colocalization of IgG and hexon in the Delta-24-RGD-treated specimens (figure 2C,D). These data indicated that intratumorally injected viruses in the brain colocalize with antibodies that may result in neutralization of the therapeutic agent. Furthermore, these results, together with the previous data showing the presence of NAbs in patients treated with Delta-24-RGD, suggest that virotherapy triggers the development of NAbs that may arrive at tumors due to the disturbance of the blood-brain barrier produced by the combination of the effect of the glioblastoma and the intracranial inflammation produced by the therapeutic agent.

**Figure 2.**
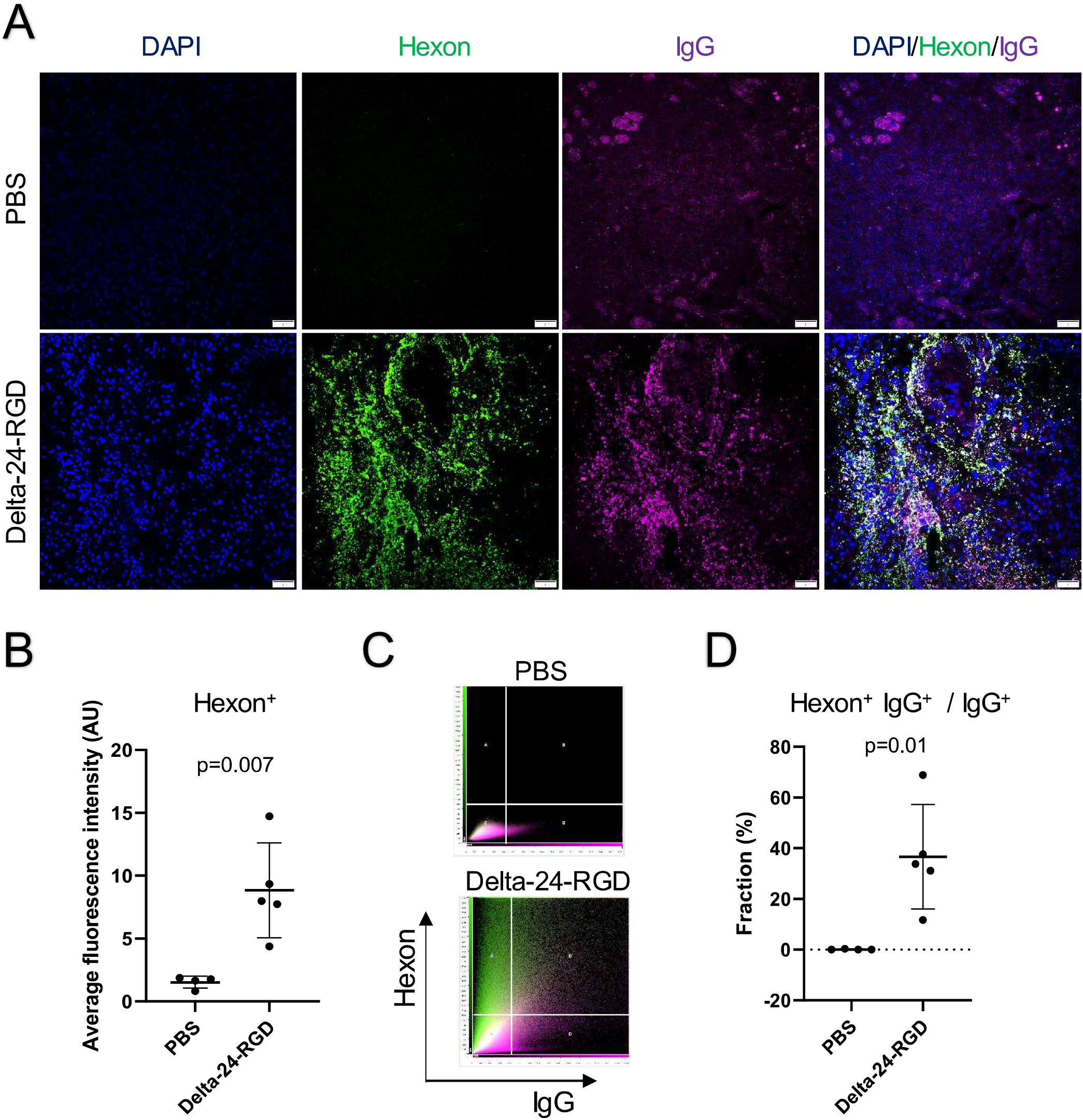
Immunoglobulins and hexon expression co-localize in Delta-24-RGD-infected murine gliomas. (A) C57BL/6 mice immunized against wildtype Ad5 were intracranially injected with GSC005 cells. After 3 intratumoral injections of PBS or Delta-24-RGD, the mice were humanely killed, and their brains were collected and processed for immunofluorescence imaging. Blue indicates DAPI; green, adenovirus hexon; and magenta, mouse IgG. Scale bar, 50 μm. (B) Quantification of adenovirus hexon fluorescence signals. Data represent mean ± standard deviation (SD) from individual mice; the *p* value was calculated using an unpaired t-test. (C) Scatter plots showing hexon and IgG signals. (D) Percentages of IgG-positive fluorescence signals in brain tumors that colocalize with hexon-positive signals. Data represent mean ± SD from individual mice; the *p* value was calculated using an unpaired t-test.

### Generation and characterization of the chimeric OV Delta-24-RGD-H43m

To overcome the challenge that NAbs might represent in virotherapy, we designed a chimeric virus called Delta-24-RGD-H43m to treat brain tumors. To this end, using molecular cloning techniques, we swapped the hexon hypervariable regions (HVRs) of Ad5, which constitutes the dominant adenovirus antigens and, therefore, the major targets for NAbs,^10,11^ with hexon HVRs from Ad43, a rare serotype to which less than 5% of the human population has been exposed and develop antibodies.^12^ Delta-24-RGD-H43m has 9 replacements in its hexon HVRs^13^ (supplementary table 1); a threonine-to-methionine substitution at position 351 (T351M; corresponding to T342M in the wildtype Ad5 hexon) that stabilizes protein folding^14^; and maintain the two mutations present in Delta-24-RGD (24–base pair deletion in E1A and RGD-4C motif insertion in the fiber) (figure 3A). A homology-based protein folding algorithm predicted that parental and chimeric hexon proteins have distinct surfaces, suggesting antibodies specific to one surface may not recognize the chimeric surface (figure 3B). Tests of thermostability of the new virus revealed that infectivity of Delta-24-RGD-H43m was not altered compared to Delta-24-RGD when the agent was exposed to several temperatures, such as 32°C, 37°C, and 42°C. Thus, DNA replications of the parental and chimeric adenoviruses were comparable at various times during the adenovirus genome replication cycle (figure 3C). We next sought to determine whether Delta-24-RGD-H43m infection produced viable and infective virus progenies. To this end, we observed that infection of human glioma U-87 MG cells, murine glioma GSC005 cells, or human lung adenocarcinoma A549 cells with either Delta-24-RGD-H43m or Delta-24-RGD induced the robust expression of both the early viral protein E1A (indicative of virus infection) and the late viral protein fiber (indicative of virus replication; figure 3D). Furthermore, transmission electron microscopy examination 72 hours after infection with either Delta-24-RGD-H43m or Delta-24-RGD identified intracellular viral progenies (figure 3E). In these microscopy images, intranuclear areas of the solid-liquid interface were in proximity to areas of adenovirus assembly, indicating active production of viral proteins and productive assembly of new virions. Together, these results suggested that Delta-24-RGD-H43m was stable and maintained similar infectious and replicative properties as Delta-24-RGD.

**Figure 3.**
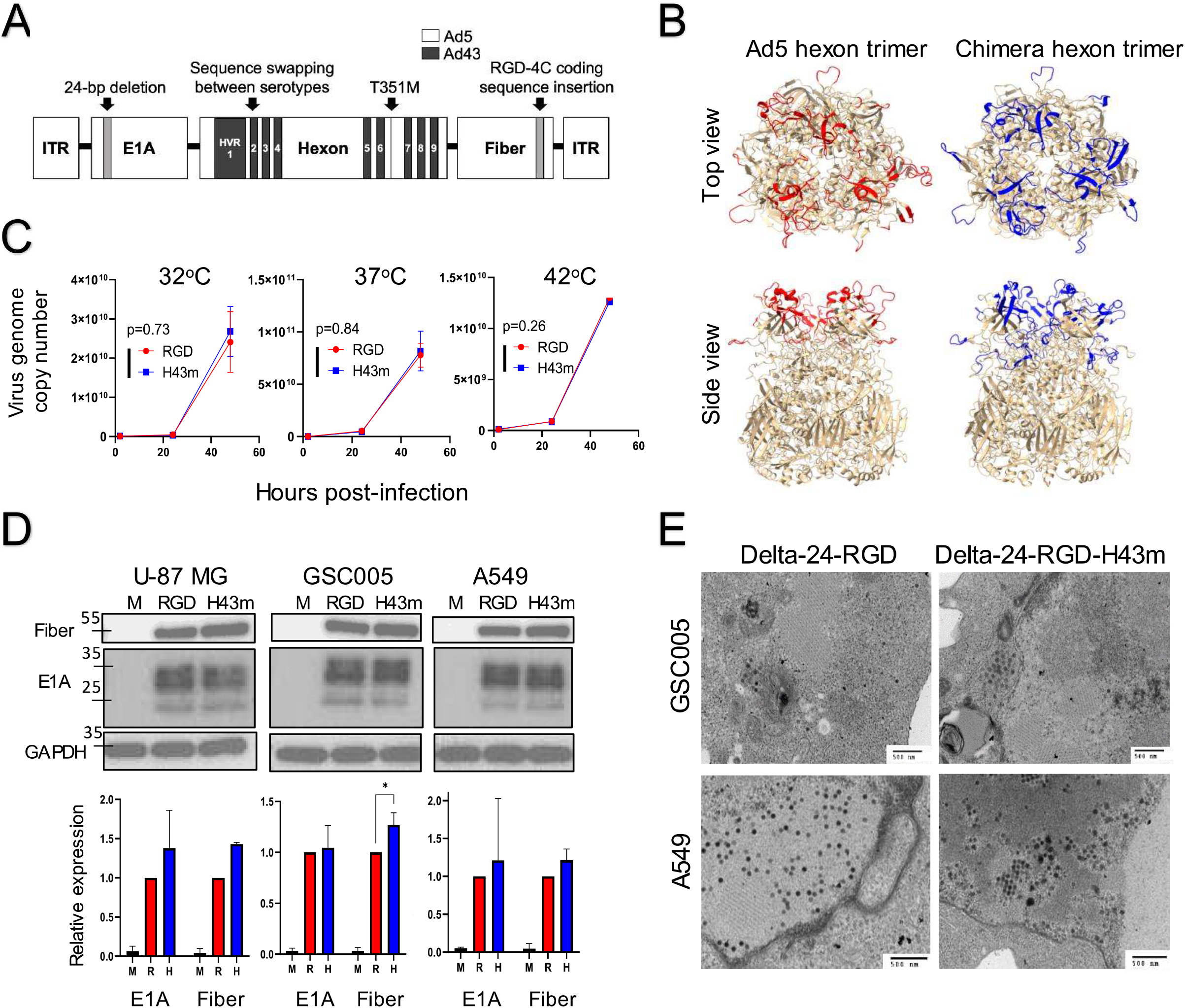
Generation and characterization of the chimeric oncolytic adenovirus Delta-24-RGD-H43m. (A) Genomic structure of Delta-24-RGD-H43m showing the mutations in the adenoviral genome, including the Ad5 and Ad43 components. (B) The predicted folded structures of the wildtype Ad5 (red) and chimeric (blue) hexon trimers were generated using the SWISS-MODEL algorithm. Hexon hypervariable regions are highlighted in colors. (C) Adenovirus genome copies were quantified using qPCR analyses of DNA extracted from A549 cells infected with Delta-24-RGD (red) or Delta-24-RGD-H43m (blue) and incubated at 32°C, 37°C or 42°C. (D) Expression of viral proteins in U-87 MG, GSC005, and A549 cells was assessed 48 hours after exposure to PBS (M) or infection with Delta-24-RGD (RGD) or Delta-24-RGD-H43m (H43m). Bar graphs show the quantification of the Western blots as mean ± SD from two to three independent experiments; M, PBS; R, Delta-24-RGD; H, Delta-24-RGD-H43m. \**P* ≤ 0.05, 2-way ANOVA. (E) Transmission electron microscopy was used to visualize adenovirus particles in GSC005 and A549 cells 72 hours after infection with Delta-24-RGD or Delta-24-RGD-H43m. Scale bar, 500 nm.

### Delta-24-RGD-H43m induces potent anti-glioma effect in vitro

Since wild-type oncolytic adenoviruses efficiently infect and lyse human and murine glioma cells, we wanted to ascertain that the chimeric oncolytic virus retains these properties in vitro. For these experiments, we plated human glioma U-87 MG, murine glioma GSC005, and human lung adenocarcinoma A549 and infected the cultures with Delta-24-RGD-H43m or Delta-24-RGD at doses ranging from 0 to 100 infectious units per cell (multiplicity of infection, MOI). Daily observation of the cells under light microscopy showed profound morphological changes and dramatic signs of cytopathic effects at doses of 40 MOIs and higher three days after the infection (data not shown). At a dose of 100 MOI, most cells infected with each of the viruses were lysed 72 hours after infection. Therefore, Delta-24-RGD-H43m infection strongly inhibited cancer cell viability in a dose-dependent manner (figure 4A). These data indicate that the modifications of the adenoviral genome that resulted in the production of Delta-24-RGD-H43m did not impair its ability to lyse cancer cells. These results are in agreement with the data obtained from thermostability and viral replication analyses.

**Figure 4.**
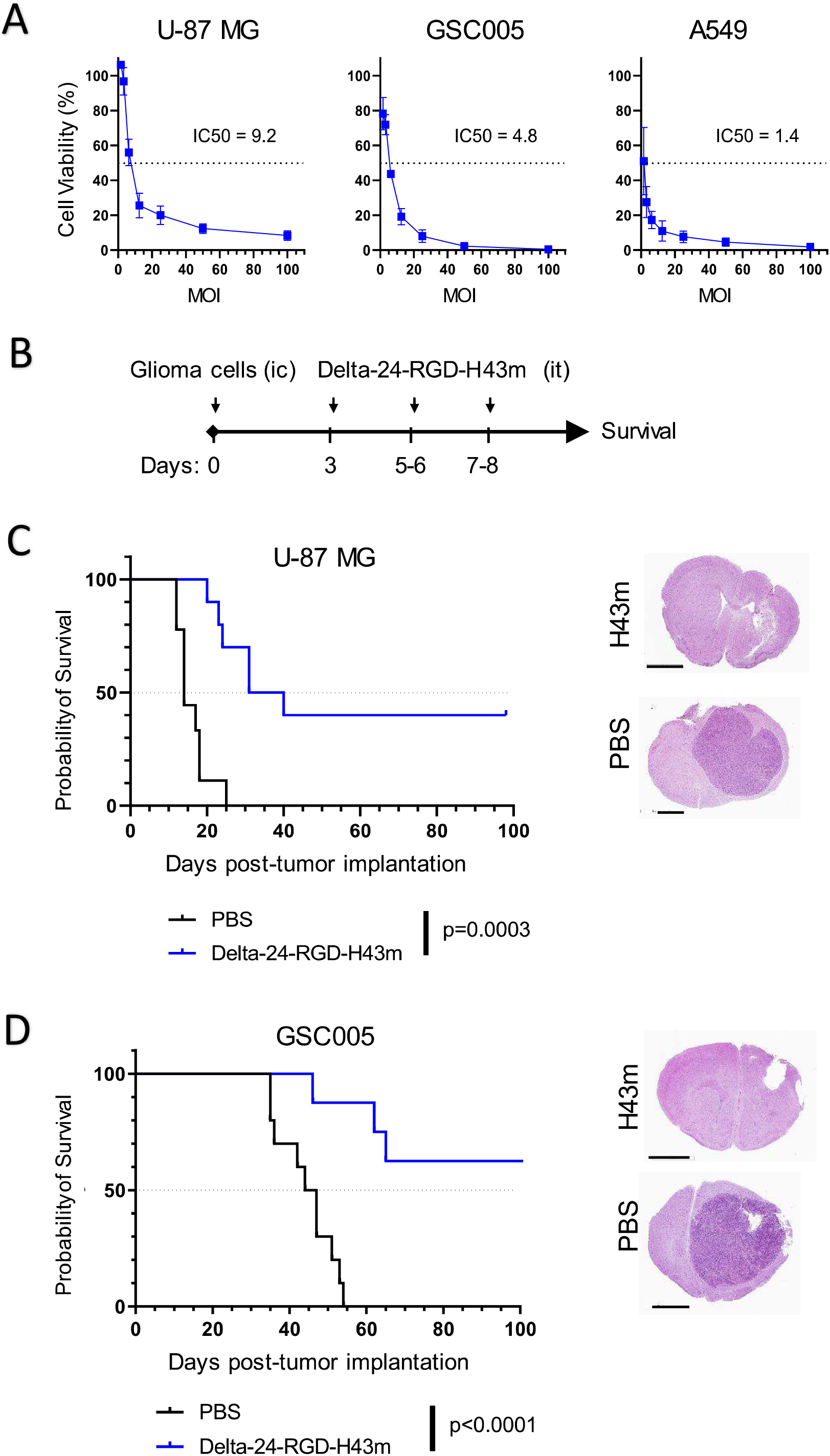
Chimeric oncolytic adenovirus Delta-24-RGD-H43m exerts potent anti-cancer effect. (A) Viability of U-87 MG, GSC005, and A549 cells were measured 72 hours after infection with Delta-24-RGD-H43m at the indicated MOIs. Data represent mean ± SD from three independent experiments. (B and C) J:NU outbred nude mice received intracranial (ic) injections of U-87 MG human glioma cells. PBS or Delta-24-RGD-H43m was injected intratumorally (it) on days 3, 5, and 7 after tumor implantation. (n=9-10 per group) (B and D) C57BL/6 mice received intracranial injections of GSC005 murine glioma cells. PBS or Delta-24-RGD-H43m was injected intratumorally on days 3, 6, and 8 after tumor implantation. n=8-10 per group) (C and D) Survival was monitored for 100 days; *p* values were calculated using a restricted mean survival time test. Brains were collected and subjected to histopathological analyses. Shown are representative images. Scale bar, 2 mm.

### Treatment with Delta-24-RGD-H43m showed robust anti-tumor efficacy in glioma-bearing mice

To determine whether Delta-24-RGD-H43m treatment leads to tumor regression, we used orthotopic models of U-87 MG human gliomas in immunodeficient J:NU nude mice and GSC005 murine gliomas in immunocompetent C57BL/6 mice (figure 4B). Compared to their mock-treated counterparts, U-87 MG glioma-bearing mice that were intratumorally treated with Delta-24-RGD-H43m prolonged median survival and resulted in long-term survivors (35.5 vs. 14 days; 40% vs 0%; *p*=0.0003, restricted mean survival time (RMST) test; figure 4C). Corroborating this result, we tested the effect of Delta-24-RGD-H43m in a syngeneic glioma model. Similarly, treatment of GSC005 glioma-bearing mice resulted in higher median survival and more long-term survivors when compared to mock treatment (undefined vs 45.5 days; 62.5% vs 0%; *p*<0.0001, RMST test; figure 4D). Postmortem examination of brains of PBS-treated mice revealed huge tumor masses at the time of their deaths. However, examination of brains in Delta-24-RGD-H43m–treated mice at the time of experiment termination (censored mice at 100 day of the experiment) showed complete regression of tumor masses (figure 4C,D). Similar results were observed in mice bearing more established intracranial GSC005 tumors, tested by initiating the treatment with intratumoral Delta-24-RGD-H43m on day 7 after tumor implantation. In this experiment, virus-treated mice had longer median survival and more long-term survivors than those that received mock treatment (61 days vs. 42 days; 11% vs 0%; *p*=0.006, RMST test; supplementary figure S2).

### Delta-24-RGD-H43m shows serotype-specific resistance to murine neutralizing antibodies

We next investigated the extent to which the chimeric virus exhibits resistance to anti-Ad5 NAbs generated in mice. We intramuscularly injected C57BL/6 mice with 2.5x10^8^ infectious units of Delta-24-RGD, Delta-24-RGD-H43m, or PBS alone. Four weeks after the first injection, the mice received a second injection to boost immunization. After the immunization process, we collected sera at week 7 (figure 5A). To test the potency of the sera to inactivate adenoviruses and calculate serum dilutions that inhibit more than 50% of virus activity, we infected A549 cells with Delta-24-RGD or Delta-24-RGD-H43m mixed with the obtained sera and quantified cell viability 48 hours later. As expected, sera from treatment-naïve mice did not inactivate the activity of either virus (figure 5B). Of interest, sera from mice immunized against Delta-24-RGD significantly inhibited Delta-24-RGD but not Delta-24-RGD-H43m (*p*=0.004, paired t-test; figure 5C). Sera from mice immunized against Delta-24-RGD-H43m showed the opposite effect and neutralized Delta-24-RGD-H43m but not Delta-24-RGD (*p*=0.03, paired t-test; figure 5D, supplementary figure S3). These results suggested that serotype-specific antibodies inhibited viral activities and confirmed that Delta-24-RGD-H43m is more resistant to anti-Ad5 antibody–mediated neutralization compared to Delta-24-RGD.

**Figure 5.**
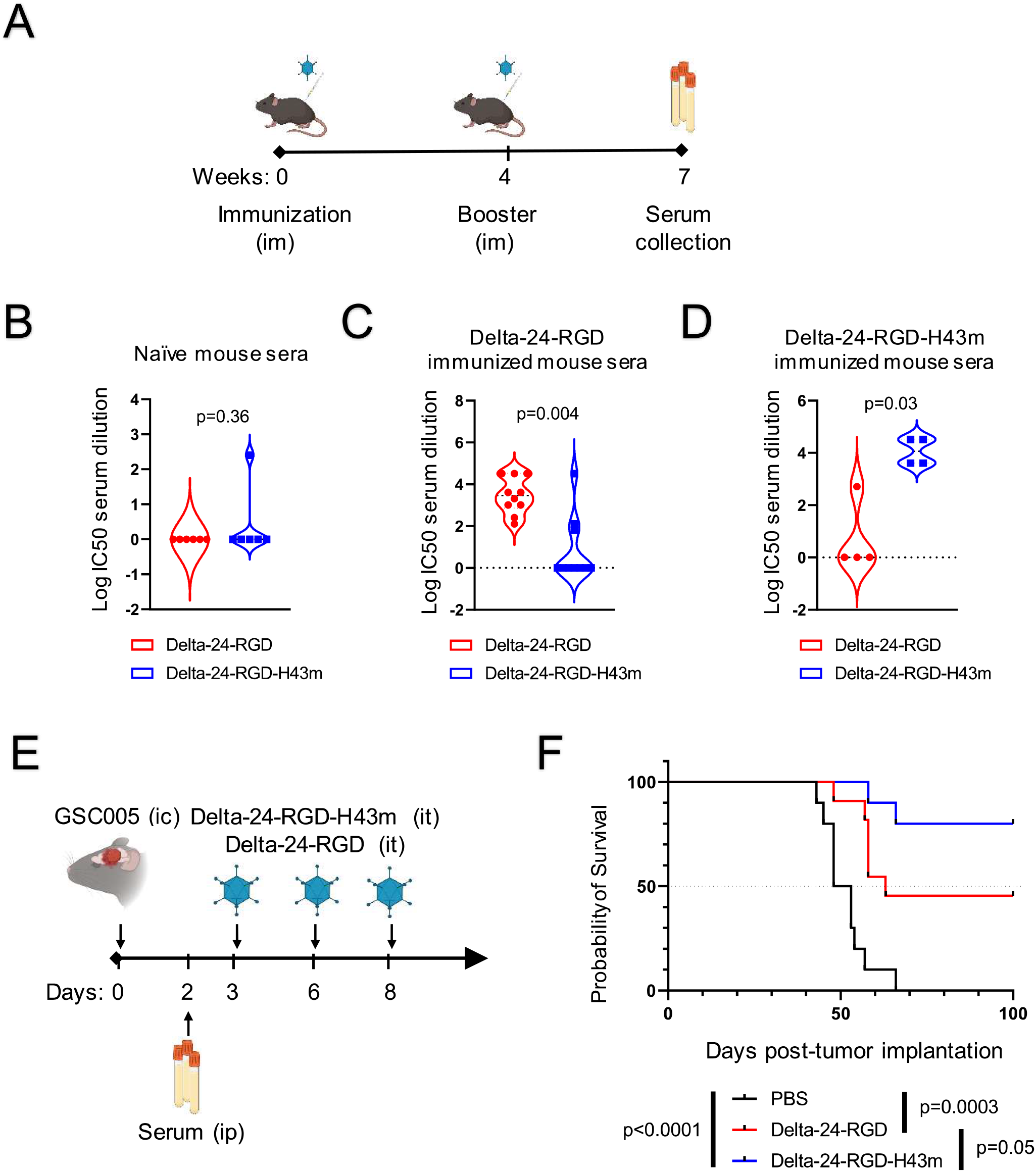
Delta-24-RGD-H43m evades anti-Ad5 NAbs from Delta-24-RGD-immunized murine model. (A) C57BL/6 mice received intramuscular injections of PBS (n=6), Delta-24-RGD (n=10), or Delta-24-RGD-H43m (n=4). Boost immunizations were performed 4 weeks later. Three weeks after boost immunization, sera were collected. (B-D) A549 cells were infected with Delta-24-RGD or Delta-24-RGD-H43m in the presence of sera collected from mice in experiment A. Cell viability was measured 48 hours later to identify the serum dilutions that inhibit 50% of cell death. Each dot represents serum from each immunized mice; *p* values were calculated using paired t-tests. (E and F). C57BL/6 mice bearing GSC005 tumors received Delta-24-RGD-H43m, Delta-24-RGD, or PBS alone injections on days 3, 6 and 8 after tumor implantation with intraperitoneal (ip) transfusions of Ad5-immunized serum on day 2. Survival was monitored for 100 days; n=10-11 per group; *p* values were calculated using restricted mean survival time tests.

### Delta-24-RGD-H43m has enhanced anti-tumor efficacy in glioma-bearing mice with anti-Ad5 immunity

To compare the therapeutic efficacies of Delta-24-RGD-H43m and Delta-24-RGD in the presence of anti-Ad5 immunity, we intracranially implanted GSC005 gliomas into C57BL/6 mice. These mice received an intraperitoneal injection of Ad5-immunized sera two days after tumor implantation, followed by intratumoral administration of Delta-24-RGD-H43m, Delta-24-RGD, or PBS alone (figure 5E). Corroborating our data from in vitro experiments, mice treated with Delta-24-RGD-H43m had longer median survival and more long-term survivors than those treated with Delta-24-RGD (undefined vs. 63 days; 80% vs. 45%; *p*=0.05, RMST test; figure 5F). Similarly, in mice bearing more established intracranial GSC005 tumors, Delta-24-RGD-H43m treatment resulted in prolonged median survival and a higher percentage of long-term survivors than those treated with Delta-24-RGD (48.5 days vs. 43 days; 10% vs. 0%; *p*=0.03, RMST test; supplementary figure S4). These data suggest that therapeutic outcomes achieved with Delta-24-RGD-H43m are substantially better than those achieved with Delta-24-RGD in animals with immunity against Ad5.

### Delta-24-RGD-H43m evades NAbs present in sera from patients treated with Delta-24-RGD

To demonstrate that Delta-24-RGD-H43m escapes the inhibitory effects of sera from patients with malignant gliomas treated with Delta-24-RGD,^5^ we infected A549 cells with Delta-24-RGD or Delta-24-RGD-H43m mixed with the obtained patient sera and quantified cell viability 48 hours later. In accordance with the results of our experiments using murine sera, the capability of Delta-24-RGD-H43m to evade the neutralization by sera collected 1 and 4 months after virus injection was significantly more potent than that of Delta-24-RGD (*p*=0.01 and *p*=0.0007, paired t-test; figure 6A,B). For example, when these viruses were mixed with sera from patient #16, the activity of Delta-24-RGD was inhibited by more than 50%, whereas the activity of Delta-24-RGD-H43m remained unchanged (*p*<0.0001, 2-way ANOVA; figure 6C). Sera from some patients, including patient #17 at 4 months, although slightly inhibited the activity of Delta-24-RGD-H43m, it was significantly more potent to inhibit Delta-24-RGD capability to lyse cancer cells (*p*<0.0001, 2-way ANOVA; figure 6C). These data strongly suggested that Delta-24-RGD-H43m would be more efficacious than Delta-24-RGD in most patients with basal anti-Ad5 immunity or that develop Nabs after an initial treatment with Ad5-based OVs.

**Figure 6.**
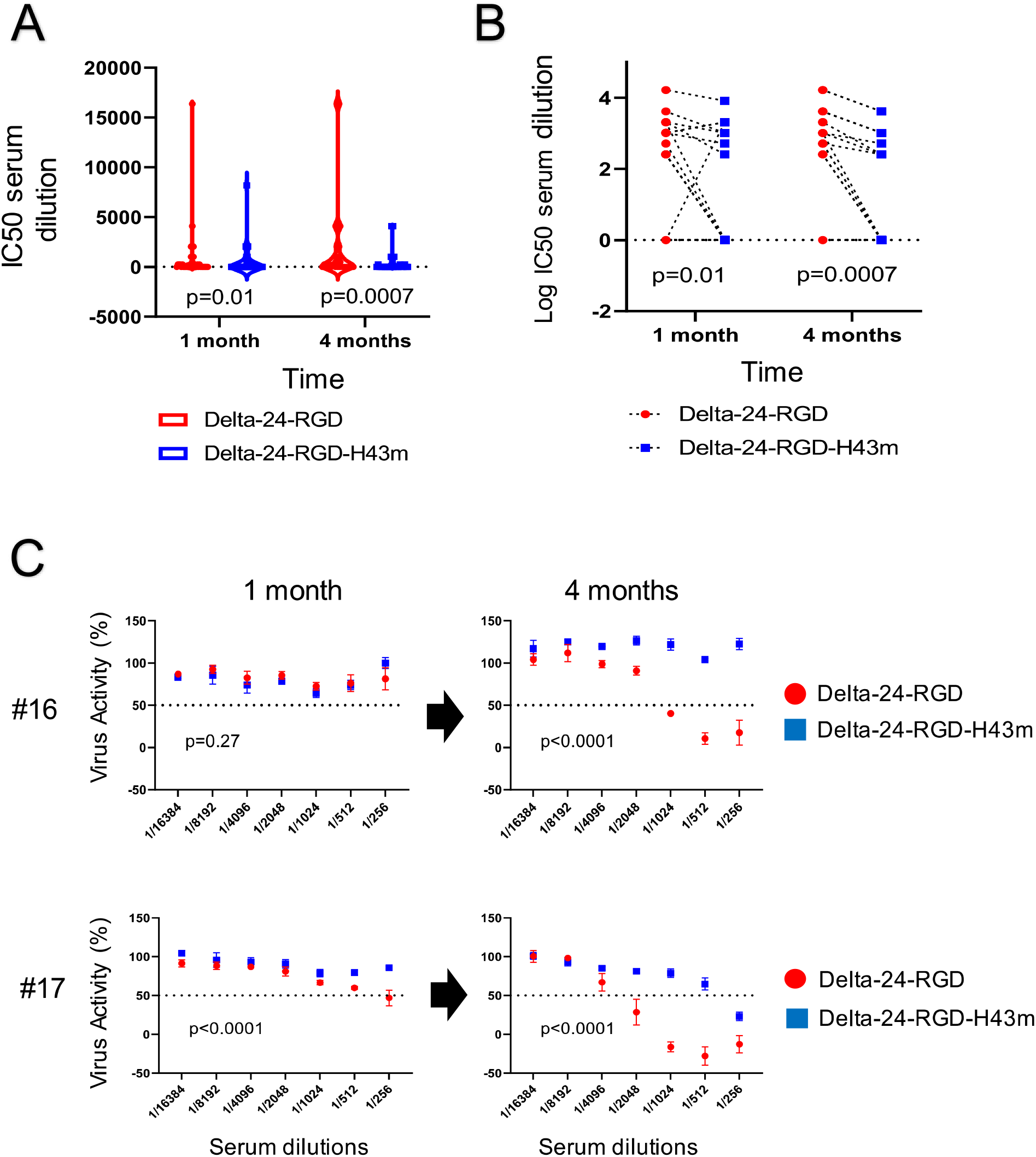
Delta-24-RGD-H43m evades NAbs from sera of patients enrolled in a phase I clinical trial of Delta-24-RGD. (A and B) Patients with recurrent high-grade gliomas (NCT00805376) were treated with Delta-24-RGD. A549 cells were infected with Delta-24-RGD or Delta-24-RGD-H43m in the presence of patient sera. Cell viability was measured 48 hours later to identify the serum dilutions that inhibit 50% of cell death. Each dot represents serum from each patient; the *p* value was calculated using a paired t-test. (C) Representative plots of virus activities across patient serum dilutions. Data represent mean ± SD at each serum dilution; *p* values were calculated using 2-way ANOVA.

## DISCUSSION

Our results highlight that adenoviruses capable of evading neutralizing antibodies against adenovirus serotype 5 should have anti-glioma efficacy in a larger percentage of patients. Furthermore, we showed that modifications of the vector backbone by replacing the adenovirus hexon immunodominant epitopes are a feasible approach to circumvent adenovirus epitope immunodominance.

Preexisting immunity to adenovirus is widely recognized as a hindrance to the widespread use of these biological agents as cancer therapies.^15^ Notably, two recent phase 1 clinical trials using oncolytic viruses to treat pediatric patients with gliomas have revealed that virus-specific antibody titers could serve as potential survival biomarkers following virotherapy.^6,16^ For instance, the median survival of pediatric patients with diffuse intrinsic pontine glioma following Delta-24-RGD treatment could be stratified according to their virus-specific NAb titers.^6^ These findings align with results from another trial involving intratumoral injection of the oncolytic virus HSV-1 G207 in pediatric patients with HGGs. In this trial, patients with baseline HSV-1 IgG antibodies had a median survival of 5.1 months, compared to 18.3 months for those without antibodies at the time of treatment.^16^ However, despite these correlations there is currently a lack of direct clinical or preclinical evidence to confirm whether antibodies within brain tumors impede viral activities.

Using an orthotopic glioma model, we first confirmed the presence of antibodies within brain tumors during virotherapy, highlighting their significance in the context of intratumoral glioma treatment. It is worth noting that previous animal studies primarily focused on examining the role of NAbs in the treatment of subcutaneous tumors, either through systemic virus injections or local intratumoral virus injections.^17–19^ Furthermore, there have been limited investigations into the impact of NAbs during virotherapy for high-grade gliomas, possibly due to the prevailing notion that the blood-brain barrier restricts antibody access to brains.^20^ Remarkably, our study is the first to reveal the presence of antibodies in brain tumors and suggest their potential interference with oncolytic virotherapy.

We found that anti-Ad5 antibodies pose a significant challenge to the effectiveness of oncolytic adenoviruses. Thus, sera from virus-immunized mice not only inhibited the cytotoxicity of Delta-24-RGD *in vitro* but also considerably limited its anti-glioma efficacy *in vivo*. These findings are in agreement with prior studies that demonstrated the inhibitory impact of Nabs on adenoviral vector activity.^15^ Of further relevance, a patient undergoing intra-arterial adenoviral vector gene therapy suffered lethal systemic inflammation caused by the formation of adenovirus-antibody complexes due to preexisting humoral immunity.^21^ These studies raise safety- and efficacy-related concerns that provided the rationale for the development of several strategies to protect therapeutic viruses from NAbs, which have included the use of polymer coatings and cellular vehicles^22,23^. Although these strategies can be useful in the case of replication-deficient viral vectors, they have limited efficacy when the therapeutic agent is a replication-competent virus because the progeny of the virus would be exposed to the immunity of the host. In the case of oncolytic viruses, genetic modifications of the virus are arguably the most feasible strategy.

The genetically modified, chimeric oncolytic adenovirus Delta-24-RGD-H43m effectively evaded anti-Ad5 NAbs, and consequently provided a significant survival benefit over Delta-24-RGD in glioma-bearing mice immunized against Ad5. However, chimeric viruses like Delta-24-RGD-H43m, which are focused on the ablation of the adenovirus dominant antigen, warrant careful consideration, as other epitopes may become targets for immunity. In this regard, other genetic modifications can be added to further diminish immunodominance. For instance, some investigators have proposed the deletion of early adenoviral genes such as E4^24^ or the replacement of adenoviral promoters^25^ to reduce Ad-specific immune responses.^26^ Although these strategies may enhance antibody evasion, they may, in turn, decrease the replication ability of the therapeutic virus.^27^ Combination of hexon and fiber modifications may also increase immune evasion.^28^ These combined strategies should be examined with detail for immune-resistant oncolytic adenoviruses and from the mechanistic angle of adenovirus antigen immunodominance. Another strategy to evade the immune response against adenoviruses would be the sequential injection of different oncolytic viruses in the same patient instead of treatment with multiple injections of the same virus. Future studies should provide further insights into the intricate interactions between different viruses and various immune factors. Nonetheless, the data from our current study, coupled with previous research, strongly buttressed the tenet that endowing oncolytic adenoviruses with the ability to partially evade antibody-mediated neutralization significantly enhances their antitumor efficacy.

In summary, analyses of our phase 1 clinical trial (NCT00805376)^5^ showed that multiple dosing of Delta-24-RGD led to the development of NAb in more than 80% of the patients, which coincided with a lack of long-term survivors. Based on the rationale derived from these clinical data, we generated and characterized a chimeric oncolytic adenovirus to evade NAbs that provided superior survival outcomes in murine models of malignant gliomas. Delta-24-RGD-H43m exhibited a robust resilience to be neutralized by patient sera, significantly improving the anti-cancer effect of the parental virus, Delta-24-RGD. Therefore, these preclinical and clinical data warrant further studies to propel a future clinical trial to explore the anti-cancer effect of genetically modified oncolytic adenoviruses designed to mitigate anti-adenovirus immunity in patients with glioma.

## MATERIALS AND METHODS

### Cell lines and culture conditions

Human lung adenocarcinoma A549 cells (ATCC, Manassas, VA) were cultured in Dulbecco’s modified Eagle’s medium–nutrient mixture F12 (DMEM/F12) supplemented with 10% fetal bovine serum (FBS) and antibiotics (100 µg/ml penicillin and 100 µg/ml streptomycin; Corning, Fort Worth, TX). Human glioma U-87 MG cells (ATCC) were cultured in minimum essential medium containing non-essential amino acid solution, 10% FBS, and antibiotics. Murine glioma GSC005 cells (kindly provided by I.M. Verma, The Salk Institute for Biological Studies, La Jolla, CA)^9^ were cultured in DMEM/F12 supplemented with N2 (Invitrogen, Carlsbad, CA), murine fibroblast growth factor-2 (20 ng/ml; PeproTech, Cranbury, NJ), murine epidermal growth factor (20 ng/ml; Promega, Madison, WI), and heparin (50 µg/ml; Sigma, St. Louis, MO). HEK293 cells (ATCC) were cultured in DMEM supplemented with 10% FBS and antibiotics.

### Generation of oncolytic adenoviruses

The construction of Delta-24-RGD was previously described.^20^ For the generation of Delta-24-RGD-H43m, a synthetic gene containing the hexon HVRs from Ad43 (replacing those in Ad5)^7,8^ was inserted into SfiI sites of the pMA-T backbone (Invitrogen). Then, a homologous DNA recombination was performed between the SfiI-cut synthetic gene and the AsiSI- and NdeI-digested pVK500C-Δ24 in *Escherichia coli* BJ5183 (Agilent Technologies, Santa Clara, CA) to produce pVK500C-Δ24-AdH43m. An additional homologous recombination between the SwaI-digested pVK500C-Δ24-AdH43m and the XbaI- and KpnI-digested pXK-F-RGD resulted in a plasmid, pAd5-D24-RGD-H43m, that contained the full oncolytic adenoviral genome with a chimeric hexon. This plasmid was digested with PacI and transfected into HEK293 cells using X-tremeGENE^TM^ HP DNA transfection reagent (Roche, Basel, Switzerland) to produce infectious viral particles. Sfil, AsiSI, NdeI, Swal, Xbal, Kpnl, and Pacl were purchased from New England BioLabs (Ipswich, MA). PCR and subsequent sequencing were used to test the integrity of the viral modifications. Oncolytic adenoviruses were amplified in A549 cells and purified using a 2-step CsCl gradient centrifugation (O.D. 260 Inc, Boise, ID). Virus titers were calculated by infecting A549 cells with serially diluted viruses and counting the hexon-expressing cells as described previously.^29^

### Immunofluorescence imaging

Nine- to 12-week-old male and female C57BL/6 mice received intramuscular injections of 2.5x10^8^ infectious units of wild-type Ad5 diluted in 50 µl of PBS in their hind limbs. Four weeks later, 50,000 GSC005 cells in 5 µl of media were intracranially implanted into the mice. Mice received intratumoral injections of PBS alone or 5x10^7^ infectious units of Delta-24-RGD diluted in 5 µl of PBS on days 3, 6, and 8 after tumor implantation. On day 8, the mice were humanely euthanized, transcardial perfusion with PBS was performed, and their brains were collected and fixed with 10% neutral buffered formalin. Brain slices were embedded in paraffin, and immunofluorescence staining was performed as described previously. Antibodies and their concentrations are given in supplementary table 2.

### Western blot analysis

Cells were seeded onto 6-well plates (300,000 cells/2 ml of media/well), and Delta-24-RGD or Delta-24-RGD-H43m was added at a concentration of 10 multiplicity of infection (MOI) for U-87 MG and A549 cells or 100 MOI for GSC005 cells. Cultures were incubated at 37°C for 48 hours, and then cells lysates were prepared using RIPA lysis buffer as described previously.^30^ Total proteins (8 µg/sample) were loaded onto 4%-20% Novex Tris-Glycine gels (Invitrogen), transferred onto polyvinylidene fluoride membrane, and detected with antibodies as described previously^24^. Antibodies and their concentrations are provided in supplementary table 2. Bands were visualized using an enhanced chemiluminescence Western blot detection system (Amersham Pharmacia Biotech, Amersham, UK).

### Cell viability assay

Cancer cells were plated in 96-well plates (10,000 cells/50 µl of media/well); 2 hours later, serially diluted Delta-24-RGD or Delta-24-RGD-H43m in 50 µl of media was added to each well. The cell-virus mixtures were incubated for 96 hours at 37°C. On day 2, 100 µl of additional media was added to the wells. On day 4, 30 µl of CellTiter-Blue® (Promega) was added to each well, and fluorescence was measured according to the manufacturer’s protocols.

### Transmission electron microscopy

A549 and GSC005 cells were seeded onto 6-well plates (300,000 cells/2 ml of media/well), and Delta-24-RGD or Delta-24-RGD-H43m were added at a concentration of 2 MOI for A549 cells or 20 MOI for GSC005 cells. Cells were incubated at 37°C for 72 hours and then fixed as described previously.^31^ The samples were polymerized in a 60°C oven for approximately 3 days. Ultrathin sections were cut in a Leica Ultracut microtome (Leica, Deerfield, IL), stained with uranyl acetate and lead citrate in a Leica EM Stainer, and examined in a JEM 1010 transmission electron microscope (JEOL, USA, Inc., Peabody, MA) at an accelerating voltage of 80 kV. Digital images were obtained using an AMT Imaging System (Advanced Microscopy Techniques Corp, Danvers, MA).

### Folding prediction modeling

DNA sequences of Ad5 hexon and chimeric hexon were used to generate prediction models using the protein structure homology-modeling server SWISS-MODEL.^32,33^

### Generation of NAb-containing mouse sera

Nine- to 12-week-old male and female C57BL/6 mice received intramuscular injections of 2.5x10^8^ infectious units of wild-type Ad5 diluted in 50 µl of PBS into their hind limbs. The mice received boost immunization 4 weeks later. Three weeks after boost immunization, blood was collected through cardiac puncture, allowed to solidify at room temperature for 30 minutes, and centrifuged at 2,000 x *g* for 10 minutes to obtain sera.

### Antibody-mediated neutralization of OVs

Sera from immunized mice or patients treated with Delta-24-RGD were heat-inactivated at 56°C for 1 hour, centrifuged at 2,000 x *g* for 10 minutes, serially diluted in media, and placed in 96-well plates (25 µl/well). Then, we added Delta-24-RGD or Delta-24-RGD-H43m at a concentration of 20 MOI in 25 µl per well. A549 cells (10,000/50 µl of media/well) were added, and the mixture was incubated at 37°C for 48 hours. On day 2, 30 µl of CellTiter-Blue® (Promega) was added to each well, and fluorescence was measured according to the manufacturer’s protocols.

### Survival studies

U-87 MG (500,000 cells/5 µl of media/mouse) were implanted into the caudate nucleus of 9- to 14-week-old male J:Nu immunodeficient nude mice (Jackson Labs) using a guide-screw system as described previously.^34^ Animals were housed in groups of 5 per cage. GSC005 cells (50,000 cells/5 µl of media/mouse) were implanted into the caudate nucleus of 9- to 14-week-old male or female C57BL/6 mice (Jackson Labs) using the same method. On indicated days, the mice were randomly assigned to the indicated control and treatment groups. The number of mice per group treatment or control is indicated for each experiment in the corresponding figure legends, decided by previous experimentation studies ^1^. Mice received intratumoral injections of 5 µl of PBS alone or with 5x10^7^ infectious units of Delta-24-RGD or Delta-24-RGD-H43m. Personnel involved in intratumoral treatments were not informed of the expected results and did not prepare the treatments or cells. For intraperitoneal serum transfusion, mice received 100 µl of immunized sera 18 hours prior to the initial virus injection. Mice were periodically monitored and euthanized when they showed signs of local or general disease or at 100 days after tumor implantation. No mice were excluded from the analyses. Brains were collected for histopathological analyses. All experimental procedures involving the use of mice were done in MD Anderson vivarium facilities, following recommended analgesia and anesthesia procedures, in accordance with protocols approved by MD Anderson’s Animal Care and Use Committee and in accordance with National Institutes of Health and United States Department of Agriculture guidelines.

### Statistical analysis

GraphPad Prism 9 was used to perform statistical analyses and generate graphs for *in vitro* and *in vivo* experiments. Two-tailed Student t-tests were used to determine statistical differences between 2 groups, and 1-way or 2-way ANOVA with multiple comparisons was used to determine statistical differences among 3 or more groups. Animal survival curves were plotted according to the Kaplan-Meier method. Survival rates of different treatment groups were compared using the restricted mean survival time analysis.

## SUPPLEMENTARY INFORMATION

This manuscript contains supplementary information.

## ACKNOWLEDGMENTS

The authors thank Dr. Inder M. Verma at The Salk Institute for Biological Studies in La Jolla, CA, for generously providing the GSC005 glioma cells; Dr. Kechen Ban and Ms. Verlene Henry for providing technical assistance with animal experiments; Mr. Kenneth Dunner, Jr. (High-Resolution Electron Microscopy Facility, MD Anderson Cancer Center) for performing the transmission electron microscopy, and Joseph Munch (Research Medical Library, MD Anderson Cancer Center) for editorial assistance.

## AUTHOR CONTRIBUTIONS

Conceptualization and design: DS, HJ, FL, MMA, CG-M, and JF. Development of methodology and acquisition of data: DS, HJ, AG, DK, XF, SS, TTN, SS, AL-R, AP, CIE, JG. Data analysis: DS, AG, TTN, and SS. Supervision: CG-M and JF. Writing, review, and revision: DS, CG-M, and JF.

## ETHICS STATEMENT

### Patient consent for publication

Not applicable

### Ethics approval

This is study involves human material and was approved by the MD Anderson Cancer Institutional Review Board for the clinical trial under number (NCT00805376). The evaluation of de-identified patient samples was approved under ID01-310. All experimental procedures involving the use of mice were done in accordance with protocols approved by the Animal Care and Use Committee of MD Anderson Cancer Center, according to National Institutes of Health and United States Department of Agriculture guidelines.

