## Supplementary material for "Chimeric oncolytic adenovirus evades neutralizing antibodies from human patients and exhibits enhanced anti-glioma efficacy in immunized mice": Supplmental Files

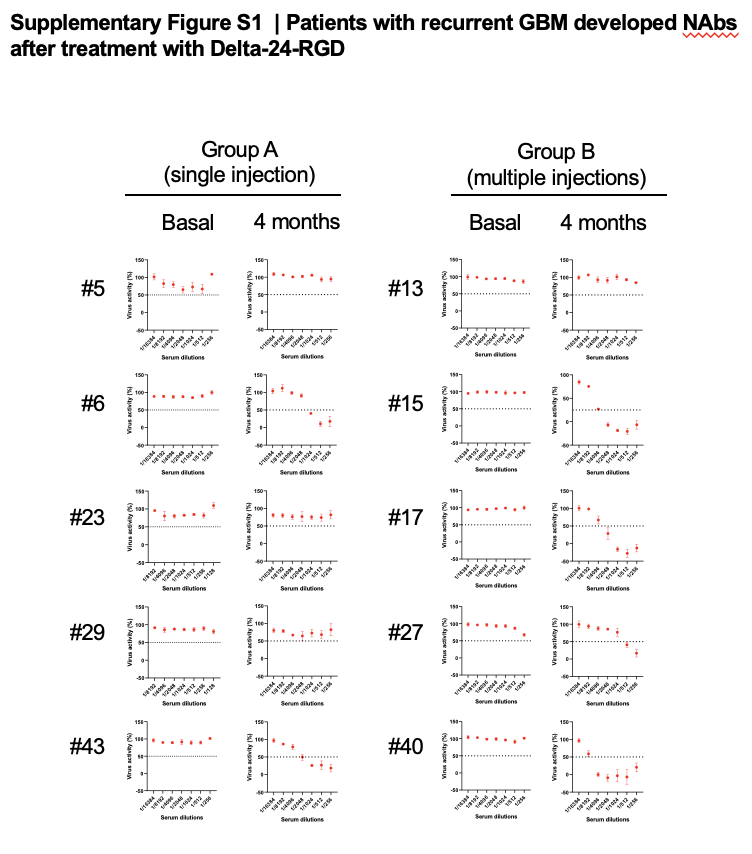

**Supplemental Figure S1. Patients with recurrent GBM developed NAbs after treatment with Delta-24-RGD.** Representative plots of Delta-24-RGD activities across serum dilutions in 10 patients with recurrent high-grade gliomas (NCT00805376) before and 4 months after initial virus treatment. Group A received a single virus injection, while group B received virus injection followed by en-bloc resection of tumors and additional virus injections into the wall of the surgical cavities. Data represent mean ± SD at each serum dilution. Dotted lines represent 50% of virus activity.

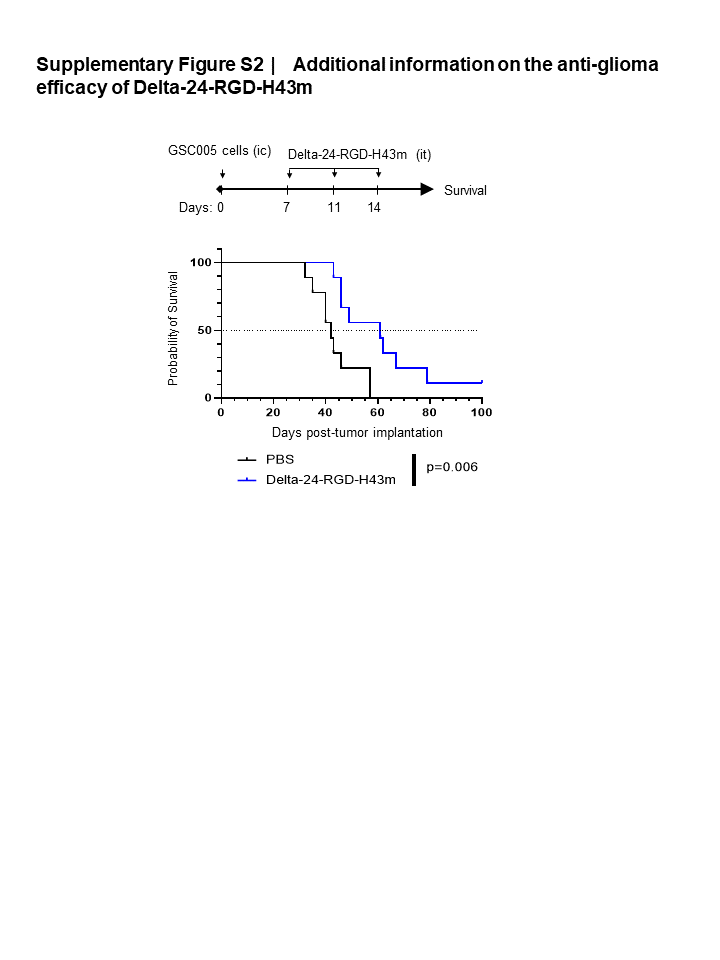

**Supplemental Figure S2. Additional information on the anti-glioma efficacy of Delta-24-RGD-H43m.** C57BL/6 mice received intracranial (ic) injections of GSC005 murine glioma cells. PBS or Delta-24-RGD-H43m (n=9 per group) was injected intratumorally (it) on days 7, 11, and 14 after tumor implantation. Survival was monitored for up to 100 days; *p* values were derived from restricted mean survival time (RMST) analysis to account for long-term survivors.

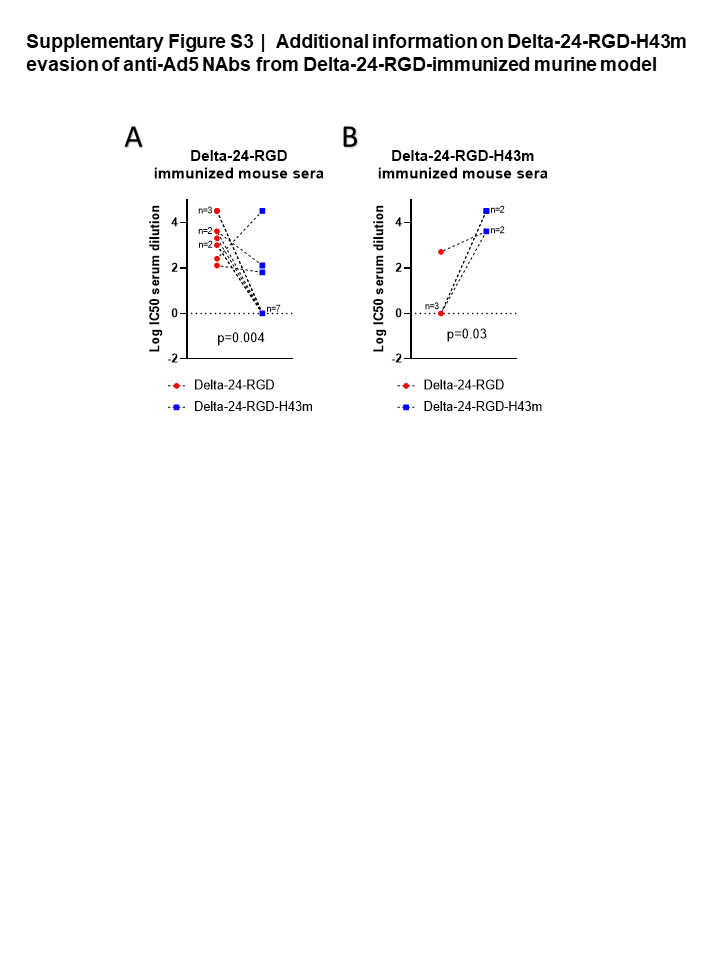

**Supplemental Figure S3. Additional information on Delta-24-RGD-H43m evasion of anti-Ad5 NAbs from Delta-24-RGD-immunized murine model.** Re-representation of Figure 5C and 5D allows comparison of log IC50 dilutions of each mouse serum for Delta-24-RGD and Delta-24-RGD-H43m viruses; *p* values were calculated using paired t-tests.

.

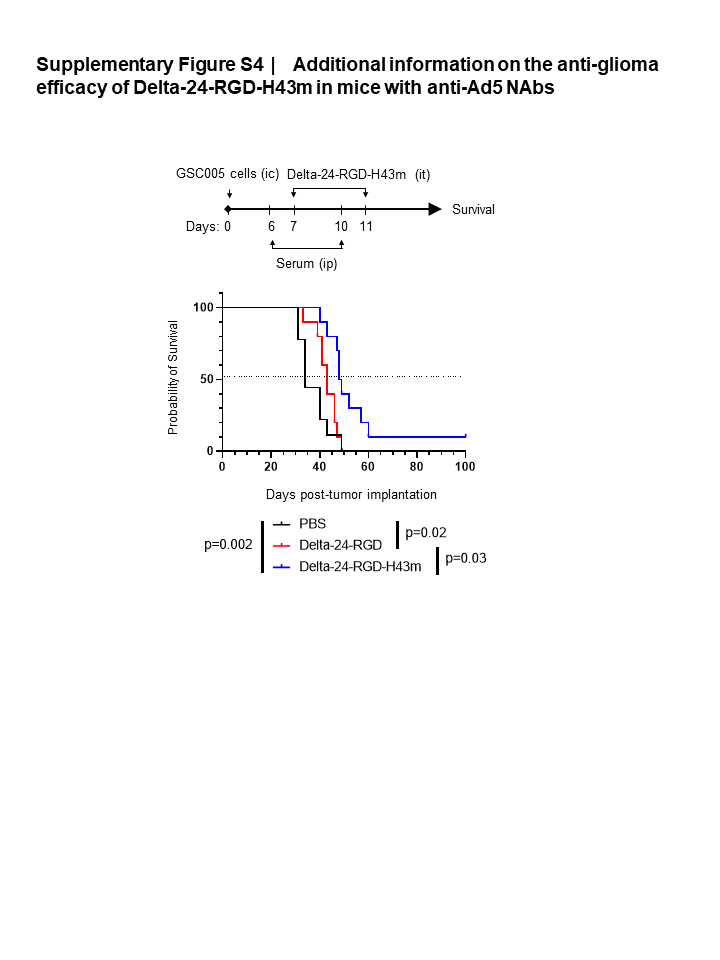

**Supplemental Figure S4. Additional information on the anti-glioma efficacy of Delta-24-RGD-H43m in mice with anti-Ad5 NAbs.** C57BL/6 mice received ic implantation of GSC005 cells. Serum collected from immunized mice (Figure 5A) were transfused into tumor-bearing mice intraperitoneally (ip) on days 6 and 10 after tumor implantation. PBS, Delta-24-RGD, or Delta-24-RGD-H43m (n=9-10 per group) was injected intratumorally 18 hours after serum injections. Survival was monitored for up to 100 days; *p* values were derived from restricted mean survival time (RMST) analysis to account for long-term survivors.
